# Effect of An Inverted Seated Position with Upper Arm Blood Flow Restriction on Measures of Elbow Flexors Neuromuscular Performance

**DOI:** 10.1101/2020.12.29.424657

**Authors:** Hamid Ahmadi, Nehara Herat, Shahab Alizadeh, Duane C Button, Urs Granacher, David G. Behm

**Author notes:** **CORRESPONDING AUTHOR:** David G Behm, School of Human Kinetics and Recreation, Memorial University of Newfoundland, St. John’s, Newfoundland and Labrador, Canada, A1C 5S7. **CONFLICT OF INTEREST:** The authors declare no conflict of interest with the contents of this manuscript.

## Abstract

**Purpose:** The objective of the investigation was to determine the concomitant effects of upper arm blood flow restriction (BFR) and inversion on elbow flexors neuromuscular responses.

**Methods:** Randomly allocated, 13 volunteers performed four conditions in a within-subject design: rest (control, 1-min upright position without BFR), control (1-min upright with BFR), 1-min inverted (without BFR), and 1-min inverted with BFR. Evoked and voluntary contractile properties, before, during and after a 30-s maximum voluntary contraction (MVC) exercise intervention were examined as well as pain scale.

**Results:** Inversion induced significant pre-exercise intervention decreases in elbow flexors MVC (21.1%, 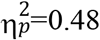, *p*=0.02) and resting evoked twitch (29.4%, 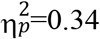, *p*=0.03) forces. The 30-s MVC induced significantly greater pre- to post-test decreases in potentiated twitch force (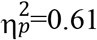, *p*<0.0009) during inversion (↓75%) than upright (↓65.3%) conditions. Overall, BFR decreased MVC force 4.8% (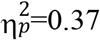, *p*=0.05). For upright position, BFR induced 21.0% reductions in M-wave amplitude (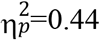, *p*=0.04). There were no significant differences for electromyographic activity or voluntary activation as measured with the interpolated twitch technique. For all conditions, there was a significant increase in pain scale between the 40-60 s intervals and post-30-s MVC (upright<inversion, and without BFR<BFR).

**Conclusion:** The concomitant application of inversion with elbow flexors BFR only amplified neuromuscular performance impairments to a small degree. Individuals who execute forceful contractions when inverted or with BFR should be cognizant that force output may be impaired.

## INTRODUCTION

Individuals can experience an involuntary inverted posture such as with an overturned vehicle or voluntary inversion with aerial maneuvers and sports (e.g. gymnastics). A decrease in force or power output, with an increased perceived difficulty in some situations can be life threatening or affect performance. Although inhibited neuromuscular function (i.e., force, rate of force development) has been reported when shifting from an upright to an inverted position (Hearn et al. 2009; Johar et al. 2013; Neary et al. 2015; Paddock et al. 2009; Smith et al. 2013), the underlying mechanisms are not clearly elucidated. Altered sympathetic nervous system activity during inversion (i.e., higher hydrostatic pressure increases vagal inputs), has been suggested as one primary mechanism that may influence changes in neuromuscular functions with inversion (Hearn et al. 2009; Johar et al. 2013; Neary et al. 2015; Paddock et al. 2009). Inversion-induced hydrostatic pressure has also been suggested to contribute to neuromuscular impairments in animals (Geeves et al. 1987; Grossman et al. 1990; Heinemann et al. 1987; Ranatunga et al. 1991) and humans (Sundberg and Kaisjer 1992).

Changes in perfusion pressure to a target muscle can be found during contractions at moderate-to-high intensities (Sadamoto et al. 1983), low-intensity contractions combined with blood flow restriction (BFR) (Yasuda et al. 2009), and when the position of a working muscle changes in respect to the level of heart (i.e., above the heart induced lower perfusion pressure) (Wright et al. 1999). Decrements in neural function and perceived exertion can be exacerbated by changes in perfusion pressure and reduced oxygen-induced peripheral fatigue (Amann et al. 2008), which for example can occur if an individual is trapped in a position with load exerted upon an immobilized limb. A squeezed or compressed limb can lead to ischaemia, increasing metabolic by-product accumulation, thereby activating pain afferents (group III and IV) (Bigland-Ritchie et al. 1986; Garland 1991; Leonard et al. 1994), contributing to central nervous system inhibition (Bigland-Ritchie et al. 1986). Decreased force production and muscle performance were observed with changes in perfusion pressure (i.e. an arm lifted above the heart level) (Fitzpatrick et al. 1996) and graded ischaemia of the lower limb (Sundberg and Kaisjer 1992). Hobbs and McCloskey (1987) indicated that with ischemia, there was greater muscle activity (electromyography: EMG) to keep the force output at the requisite level. Partial occlusion (increased pressure and ischaemia) concomitant with prolonged exercise could influence the perception of effort and sense of pain (Hollander et al. 2010). Although both inversion (Hearn et al. 2009; Johar et al. 2013; Neary et al. 2015; Paddock et al. 2009) and blood flow restriction (Amann et al. 2008, Fitzpatrick et al. 1996, Hobbs and McCloskey 1987, Sundberg and Kaisjer 1992) can alter force output and muscle activation, the combination of inversion (increased hydrostatic pressure) with blood flow restriction (BFR: increased perfusion pressure), on neuromuscular performance has not been previously investigated. It is unknown whether additive effects of hydrostatic (whole body effects) and perfusion (local muscle effects) pressure occur (inversion and BFR respectively), exacerbating neuromuscular performance decrements.

With these contexts, the objective of this study was to investigate the potential effects of a one-minute inverted position with upper arm BFR on measures of elbow flexors isometric maximum voluntary contraction (MVC) force production, biceps and triceps brachii electromyographic (EMG) activity, perceived pain, and performance fatiguability (extent of force reduction with a 30-s MVC). Based on prior inversion studies (Hearn et al. 2009; Johar et al. 2013; Neary et al. 2015; Paddock and Behm 2009), it was hypothesized, that inversion and BFR would decrease both voluntary and evoked force output, decrease voluntary activation, and increase fatigue during the 30-s MVC, and the addition of BFR to inversion would amplify these impairments to neuromuscular function.

## METHODOLOGY

### Participants

Based on the force data from previous studies on similar research topics (Hearn et al. 2009; Johar et al. 2013), a statistical a priori power analysis (G*Power 3.1, Dusseldorf, Germany) indicated that a minimum of six participants would be needed to attain an alpha of 0.05 (α error) with an actual power of 0.8 (1-β error). We were able to recruit a convenience sample of 13 (males, n=7, age: 24.7 ± 4.9, height: 178.3 ± 8.3 cm, and mass: 79.9 ± 8.6 kg and females, n=6, age: 24.5 ± 4.8, height: 162.0 ± 3.6 cm, and mass: 73.5 ± 14.3 kg) healthy physically active university students. Participants performed structured physical activity 3-4 days per week and had no previous history of cerebral, hypertensive, or visual health problems or injuries. The participants were given an overview of all procedures (i.e., orientation and testing sessions) before data collection. If willing to participate, participants signed the consent form and completed a ‘Physical Activity Readiness Questionnaire for Everyone’ (PAR-Q+: Canadian Society for Exercise Physiology, approved September 12, 2011 version). The Interdisciplinary Committee on Ethics in Human Research, Memorial University of Newfoundland (ICEHR Approval #: 20192154-HK) approved this study and the study was conducted in accordance with the latest version of the Declaration of Helsinki.

### Experimental design

Based on the recommendation of the Canadian Society for Exercise Physiology (Health Canada 2004), participants were advised to not smoke, drink alcohol or partake in intensive physical activity six hours prior to testing and to not eat food two hours before participating in the testing procedure. Participants attended an orientation session, at least a week before data collection, where they were familiarized with both upright and inversion postures; and also became familiar with BFR, the interpolated twitch technique (ITT), EMG electrode placement, and isometric maximal voluntary contractions (MVC) techniques. Following the orientation session, all participants were capable of achieving near maximal or maximal voluntary activation before commencing the experimental conditions. Using random allocation (generated by Microsoft Excel) participants performed four, ∼60 minutes, experimental conditions in a within subject design: 1) Control (1-min upright position without BFR), 2) Control with BFR (1-min upright position with BFR), 3) 1-min inversion (without BFR), and 4) Combined: 1-min inversion with BFR. Testing included 1) initial testing (upright seated position pre-fatigue), 2) pre-testing (one of the four conditions pre-fatigue) and post-testing after the 30-s MVC (one of the four conditions after the 30-s MVC). There was approximately 48 h between each experimental condition. To decrease diurnal rhythms effects, all of the test procedures were completed at approximately the same time of day. The temperature of the laboratory was maintained at 20C.

Maximal evoked muscle twitch (i.e., resting position) and voluntary (MVC with ITT) contractile properties were initially tested from an upright seated position. The participant then performed a warm-up, which consisted of arm cycling (cycle ergometer: Monark Ltd. Sweden), at 70 rpm with 1 kp for 5-min; followed by a specific warm-up of five 5-s isometric elbow flexion (∼50% of perceived maximum) contractions. Following the warm-up and a 5-min rest period, pre-fatigue BFR testing procedures (relative to the posture) with the participant positioned in the inversion chair were assessed. The subsequent condition/position (without/with an upper right arm BFR) also included an evoked twitch (10-s after achieving desired position or condition), MVC with ITT (5-s after the evoked twitch), a 30-s MVC protocol (5-s after MVC), followed by another evoked twitch and MVC with ITT (5- and 10-s after 30-s MVC respectively) (Figure 1). It is noted that there was a 5 s transition from both upright to supine, and supine to inversion positions. In total, a participant was placed at each condition for 150s, from positioning at the desired posture to post-30s MVC measures (i.e., post-fatigue evoked twitch, MVC ITT, and potentiated twitch force).

**Figure 1:**
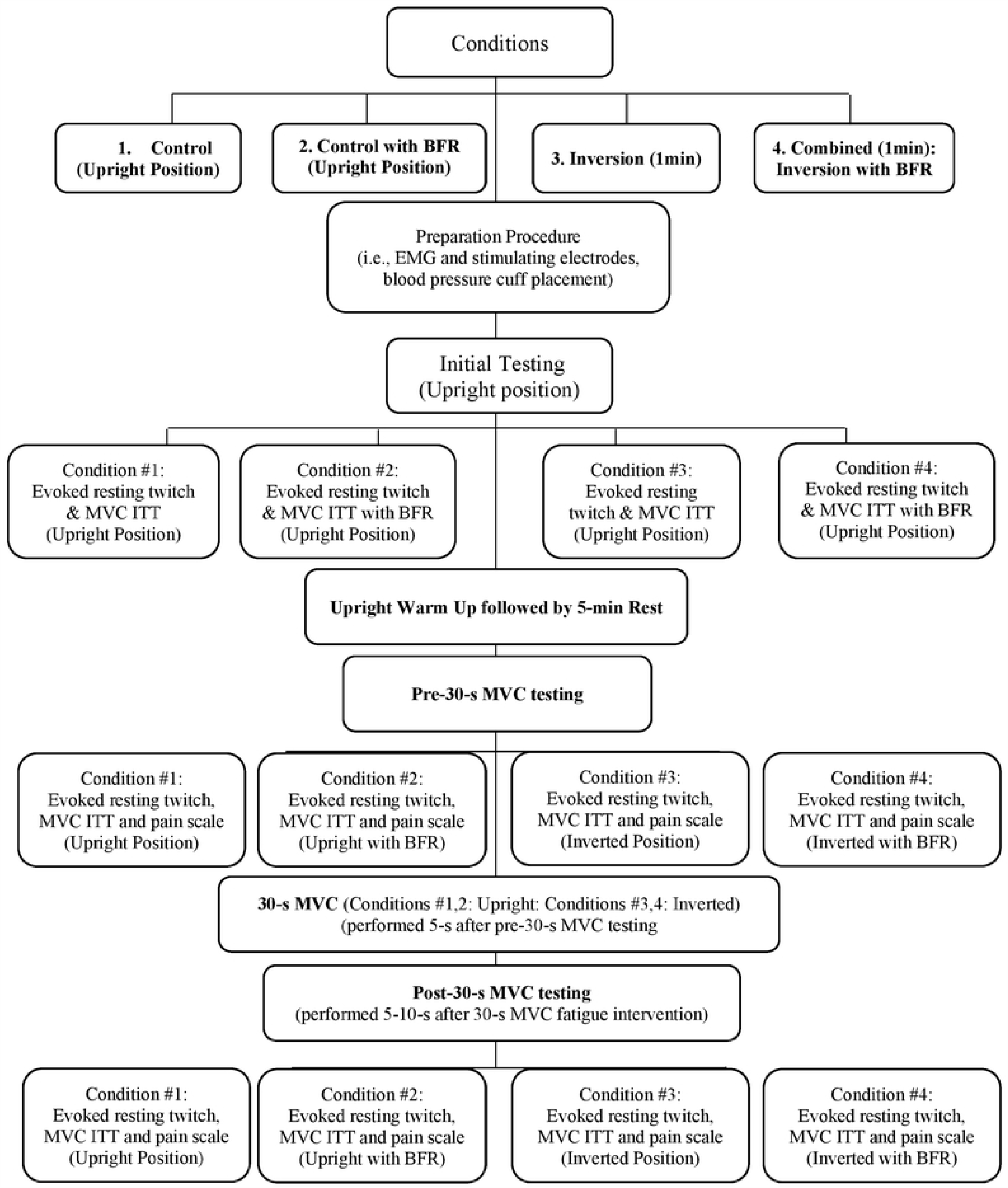
Experimental Design Acronyms: BFR: blood flow restriction, EMG: electromyography, ITT: interpolated twitch technique, MVC: maximal voluntary contraction

### Force measures

For voluntary and evoked force measures, the participants were seated in an inversion chair (initially in an upright position), which was designed and constructed by Technical Services of Memorial University of Newfoundland (Hearn et al. 2009; Neary et al. 2015; Paddock and Behm 2009)(Figure 2). The chair can rotate through a 360-degree range. Straps secured the participant at the head, torso, shoulder, hip, and thighs. Hips and knees were positioned at 90° during data collection. A Wheatstone bridge configuration strain gauge (Omega Engineering Inc., Don Mills, Ont.) via a high-tension wire cable was attached to a reinforced strap around the right wrist to assess force output, and forces were collected and amplified via analog to digital data collection hardware and software (i.e., Biopac System Inc. DA 100, A/D convertor MP100WSW; Holliston, MA). Due to the physical layout of the laboratory equipment, the right elbow flexors were tested.

**Figure 2:**
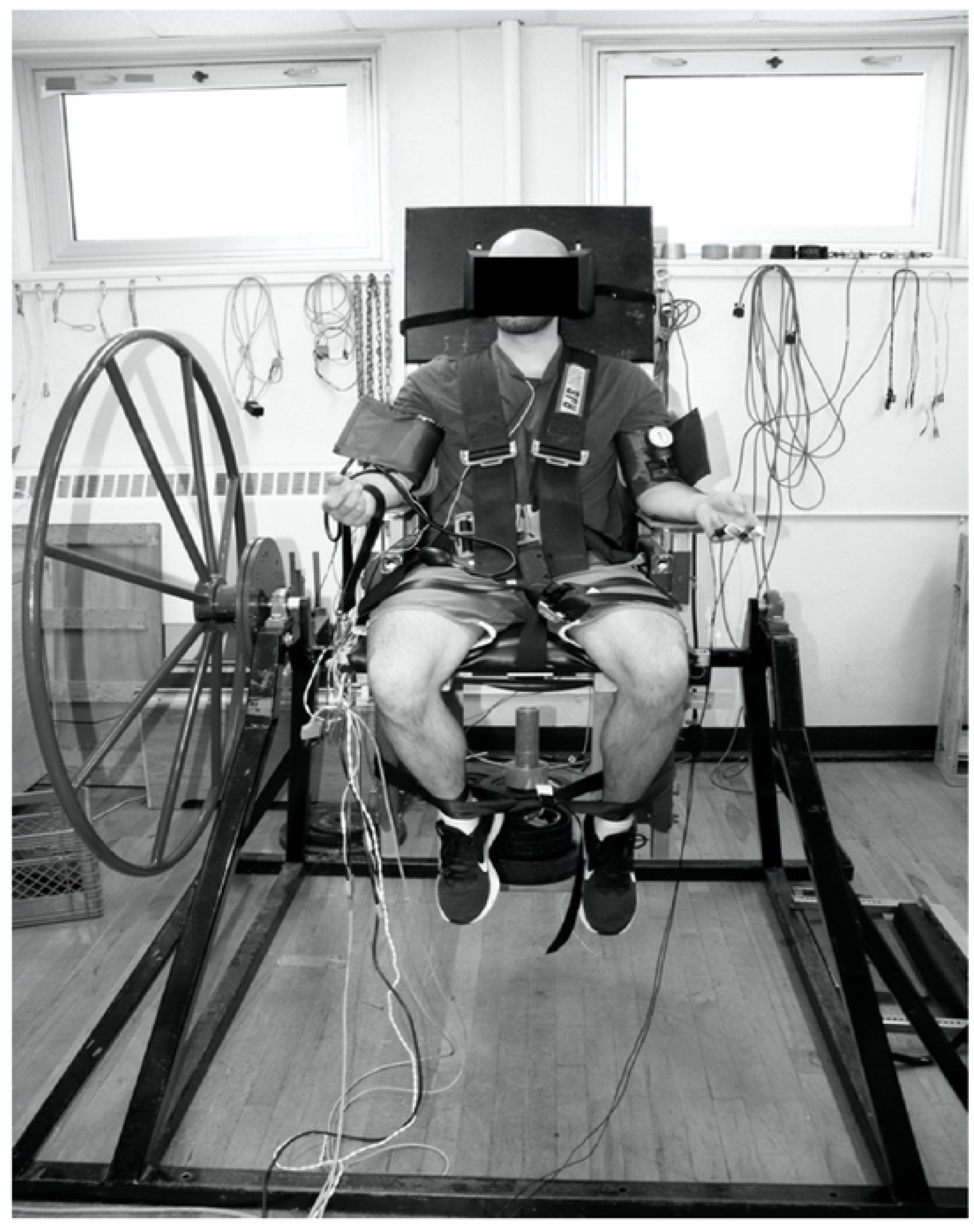
A seated participant within the inversion chair (upright posture)

### Evoked contractile properties

To assess evoked twitch forces and muscle compound action potentials (M-wave), evoked contractile properties were measured. To stimulate the musculocutaneous nerve, stimulating electrodes (electrode width was 5 cm) were wrapped around the intersection of the biceps brachii and deltoid (anode) and antecubital space (cathode), respectively (Halperin et al. 2014). Furthermore, the placement of electrodes was marked with ink from test to test to maintain the correct position of electrodes during each session. Stimulating electrodes were connected to a stimulator (Digitimer Stimulator, Model DS7AH, Hertfordshire, UK) with a maximum of one ampere (A) and 400 volts (V). Then, both amperage and voltage were increased sequentially until a plateau in the twitch torque was attained. While the initial resting twitch involved a single stimulus, the superimposed and subsequent potentiated twitches had a 10 ms inter-pulse interval between two maximal twitches during biceps brachii nerve stimulation (Behm et al. 1996). The ITT has been reported to be a valid and reliable measure of muscle inactivation (Behm et al. 1996).

Previous studies (Behm and St-Pierre 1997a;b; Behm et al. 1996; Merton 1954) have shown that there is a possibility to activate all muscle fibres via a superimposed twitch on a voluntary contraction (ITT). For consistency, each session commenced with an initial evoked twitch. Since evoked single twitches are sensitive to prior contractions resulting in a potentiated response, the twitches were evoked both prior to (resting twitch) and after (potentiated twitch) the MVC testing (Behm 2004; Behm et al. 2004). Superimposed twitches were delivered during 2-3 MVCs (4-s duration with 2-min rest between each MVC), before the BFR and fatigue intervention, and once following the BFR and 30-s MVC fatiguing muscle contraction (i.e., MVC during elbow flexion). The first supramaximal (120% of maximum) electrical stimulation was delivered at the 3-s point of the MVC (all participants could achieve maximal force within this duration), and the second twitch (as a potentiated twitch) was evoked at a 3-s interval after the MVC (subject was instructed to relax). This procedure was repeated if the MVC force of the second MVC was 5% higher than the first MVC (Behm et al. 2004; Behm and St-Pierre 1997a;b; Behm et al. 1996; Button and Behm, 2008; Halperin et al. 2014; Hearn et al. 2009). To estimate maximal voluntary activation, the amplitudes of the superimposed and post-contraction stimulation were compared [voluntary activation = [1 – (superimposed twitch/potentiated twitch)] * 100] (Behm et al. 1996).

A fatiguing protocol consisting of a 30-s MVC of the elbow flexors, was performed using an isometric elbow flexion MVC (right arm). The researcher provided consistent verbal encouragement in terms of wording and timing (e.g., “keep it up” every 10 s, starting at 10 s point).

### Voluntary contractile properties

With the same set-up as the evoked contractile properties, elbow flexors MVC isometric force was measured while seated and secured in the inversion chair with a supinated forearm at a 90° elbow flexion angle with shoulders at 0° (mid-frontal plane), with a reinforced strap around the right wrist. An instruction (“as hard and as fast as possible”), with verbal encouragement (“go go”) was provided by the researcher during the entire 4-s isometric MVC. The forces detected by the strain gauge, were used to analyze the peak isometric MVC.

### Electromyography (EMG)

Surface EMG was monitored during evoked twitches (muscle action potential: M-wave), 4-s MVC and 30-s MVC. First, the skin was prepared before electrode placement with shaving (removal hair), abrading (to remove possible dead epithelial cells), and cleaning the area with an alcohol swab to remove oils. Then, two pairs of electrodes (Kendall Medi-trace 100 series, Chikopee, Mass.) were positioned (established by SENIAM, Hermens et al. 1999) at the mid-belly of the biceps brachii (i.e., at 50% on the line from medial acromion to the fossa cubit) as an agonist muscle (corresponding to the muscle fibres), and the lateral head of the triceps brachii (i.e., halfway from the posterior crista of the acromion to the olecranon as an antagonist muscle). The reference electrode was positioned on the ulnar styloid process.

The EMG signal was collected at 2000 Hz, band-pass filter 10-500 Hz and amplified 1000x (Biopac System MEC 100 amplifier, Santa Barbara, Calif; input impedance = 2M, common mode rejection ratio > 100 dB minimum [50/60 Hz]). The collected data (via the A/D converter, Biopac MP150) was stored on a personal computer for post-processing analysis. The final data (i.e., raw EMG) was rectified and integrated over 500 ms following an MVC (Johar et al. 2013; Neary et al. 2015; Paddock and Behm 2009).

### Blood flow restriction (BFR)

To decrease potential harmful biological effects, which can occur following an extremely high dose of BFR (Loenneke et al. 2014), previous evidence (Takarada et al. 2000) using Doppler ultrasonograms showed that a moderate BFR and partial occlusion of the brachial artery can lead to 100% venous restriction and partial arterial BFR, respectively. In order to reduce hydrostatic pressure effects, and perfusion to the right elbow flexors muscles, BFR was implemented. At the resting position, a pressure cuff (A+ Med 7-62 pressure cuff; Toronto, Canada) was placed around the upper right arm, while positioned at the level of the heart. Then, the hand bulb was squeezed to manually find a pulse elimination pressure (100% BFR). The individualized BFR in the present study was set relative to brachial systolic blood pressure. To create partial BFR for upright/inverted position, the hand bulb was squeezed to 50% of complete BFR. Furthermore, with a maximal isometric contraction, the blood flow would be further restricted, obviating the need for maximal BFR with a cuff.

### Universal pain assessment tool

The researcher utilized a universal pain assessment tool (scale with numbers and cartoon facial figures)(Belcheva and Shindova 2014) to assess the degree of overall discomfort (pain perception) during inversion. Also, to investigate whether there was any interaction between the effects of inversion and BFR (during two control sessions), the sense of discomfort with and without BFR was examined.

### Data analysis

With the single electrical stimulation, peak twitch force, time to peak twitch force, half relaxation time and M wave were measured for resting and potentiated twitches. Peak MVC forces (Perrine and Edgerton 1978) were analyzed. The resting force output in the upright position was zeroed and used as the baseline when compensating for the suspended arm mass in the inverted position. The mass of the arm was recorded in the inverted position and this value subtracted from the peak force in order to counterbalance the force of gravity negatively affecting force output in the upright position. A force fatigue index was calculated from the 30-s MVC, which involved dividing the mean force of the last 5-seconds of the fatigue protocol into the mean force output of the first 5-seconds.

The integrated EMGs, from the biceps and triceps brachii, were analyzed from a one second period of the 3-s MVC (before the superimposed twitch) (Button and Behm 2008). Furthermore, a Fast Fourier transform was used to report the EMG median frequency following the fatigue protocol, as it is a reliable indicator for signal conduction velocity (following a fatigued-exercise) and generally considered a more sensitive indicator of fatigue than a raw EMG signal (Agarwal and Gottlieb 1975; Daanen et al. 1990; Dimitrova and Dimitrov 2003; Kwatny et al. 1970).

### Statistical analysis

The SPSS software (version 23.0, SPSS, Inc. Chicago, IL) was used for statistical analysis. Normality of the data was assessed and confirmed using a histogram chart to illustrate skewness and kurtosis with a Shapiro-Wilks test. The value of Greenhouse-Geisser were reported if the assumption of sphericity was not met. Initially, sex effects were considered as a factor, but with a lack of any significant differences, the sample population was integrated for analysis. To determine the effect of inversion with BFR on fatigue index of the 30-s MVC, a two-way repeated measures ANOVA (2 seated positions × 2 blood flow conditions), while three-way repeated measures ANOVA (2 seated positions × 2 blood flow conditions × 2 times) was conducted for EMG median frequency. For the evoked twitches, MVCs, ITT, potentiation twitches, muscle action potential (M-) waves, EMG, a three-way ANOVA (2 seated positions and 2 blood flow conditions and three times [initial upright resting position, pre-, and post-30-s MVC]) was applied. Further, a repeated measures ANOVA was conducted to analyze the pain scale. Differences were considered significant when a minimum value of *p*=0.05 was reached. Planned pairwise comparisons; Bonferroni adjustment, was selected to compare main effects. Additionally, the calculated partial eta squared 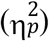 by SPSS was reported as a magnitude of outcomes (effect sizes); which is classified as small 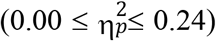, medium 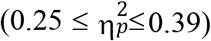, and large 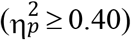 (Cohen 1998). Day to day reliability of measures (for initial upright resting position test) was assessed with Cronbach’s alpha intraclass correlation coefficient (ICC).

## RESULTS

Within the results text and figures, significant interactions are reported. Significant main effects are listed and described in Table 1.

**Table 1.** Main Effects (means ± standard deviation)

| Main Effects for Time | Initial Test | Pre-Fatigue | Post-Fatigue | Initial to Pre-Fatigue | Pre to Post-Fatigue |
| --- | --- | --- | --- | --- | --- |
| MVC Force (N)<br>$F_{(2,16)} = 52.58$ | 309.32 ± 84.55 | 269.18 ± 75.16 | 226.30 ± 63.38 | $p < 0.001$, $\eta_p^2 = 0.82$,<br>$\Delta 14.9\%$ | $p < 0.001$, $\eta_p^2 = 0.84$<br>$\Delta 18.9\%$ |
| F100 (N)<br>$(F_{(2,18)} = 31.81)$ | 73.28 ± 36.47 | 52.14 ± 25.64 | 34.32 ± 18.17 | $p = 0.003$, $\eta_p^2 = 0.63$<br>$\Delta 40.5\%$ | $p < 0.001$, $\eta_p^2 = 0.80$<br>$\Delta 51.9\%$ |
| BB EMG (mV/s)<br>$F_{(1,12)} = 22.41$ | 0.30 ± 0.16 | 0.29 ± 0.15 | 0.24 ± 0.12 | $p > 0.05$ | $p < 0.001$, $\eta_p^2 = 0.65$<br>$\Delta 22.0\%$ |
| TB EMG (mV/s)<br>$F_{(1,8)} = 7.34$ | 0.07 ± 0.02 | 0.06 ± 0.02 | 0.05 ± 0.01 | $p = 0.02$, $\eta_p^2 = 0.47$<br>$\Delta 16.6\%$ | $p = 0.003$, $\eta_p^2 = 0.67$<br>$\Delta 20.0\%$ |
| BB M-wave (mV)<br>$F_{(1,7)} = 3.89$ | 9.05 ± 4.06 | 8.96 ± 3.22 | 8.49 ± 2.95 | $p > 0.05$ | $p = 0.08$, $\eta_p^2 = 0.35$<br>$\Delta 7.6\%$ |
| PTF (N)<br>$F_{(2,10)} = 36.3$ | 66.48 ± 19.11 | 63.82 ± 18.38 | 28.86 ± 14.07 | $p > 0.05$ | $p < 0.001$,<br>$\eta_p^2 = 0.87$<br>$\Delta 121.1\%$ |
| VMA (%)<br>$F_{(1,4)} = 6.62$ | 91.71 ± 5.36 | 92.47 ± 3.44 | 82.53 ± 11.35 | $p > 0.05$ | $p = 0.06$, $\eta_p^2 = 0.62$<br>$\Delta 12.0\%$ |
| Main Effects for BFR | no BFR Conditions |  | BFR Conditions | no-BFR to BFR |  |
| MVC Force (N)<br>$F_{(1,8)} = 4.84$ | 274.61 ± 76.60 | | 261.93 ± 72.12 | $p = 0.05$, $\eta_p^2 = 0.37$<br>$\Delta 4.8\%$ | |
| BB M-Wave (mV)<br>$F_{(1,7)} = 7.28$ | 9.67 ± 2.11 | | 7.99 ± 2.44 | $p = 0.03$, $\eta_p^2 = 0.51$<br>$\Delta 21.0\%$ | |
| Perceived Pain Scale<br>$F_{(1,12)} = 7.85$ | 2.5 ± 2.59 | | 3.19 ± 2.27 | $p = 0.01$, $\eta_p^2 = 0.39$<br>$\Delta 15.6\%$ | |
| Main Effects for Seated Position | Upright Seated Positions |  | Inverted Seated Positions | Upright vs. Inverted |  |
| F100 (N)<br>$F_{(1,9)} = 3.42$ | 58.27 ± 28.01 | | 48.22 ± 25.51 | $p = 0.09$, $\eta_p^2 = 0.27$<br>$\Delta 8.3\%$ | |
| Perceived Pain Scale<br>$F_{(1,12)} = 14.56$ | 2.26 ± 1.98 | | 3.42 ± 2.8 | $p = 0.002$, $\eta_p^2 = 0.54$<br>$\Delta 26.5\%$ | |
| Main Time Effects for Fatigue-Force, and Fatigue-EMG Relationships | 0-5s interval (no-BFR and BFR) |  | 25-30s interval (no-BFR and BFR) | 0-5s to 25-30s intervals |  |
| Force (N)<br>$F_{(1,11)} = 29.96$ | 278.42 ± 85.72 | | 242.86 ± 74.45 | $p < 0.001$, $\eta_p^2 = 0.73$<br>$\Delta 14.6\%$ | |
| BB EMG Median Frequency (Hz)<br>$F_{(1,10)} = 84.65$ | 57.67 ± 10.61 | | 41.32 ± 5.59 | $p < 0.001$, $\eta_p^2 = 0.89$<br>$\Delta 39.5\%$ | |
| TB EMG Median Frequency (Hz)<br>$F_{(1,12)} = 86.69$ | 59.25 ± 12.53 | | 44.26 ± 9.25 | $p < 0.001$, $\eta_p^2 = 0.87$<br>$\Delta 33.8\%$ | |

\*BB = Biceps Brachii, EMG: electromyography, F100: force output within the first 100 ms of the MVC, MVC: isometric maximal voluntary contraction, M-wave: muscle action potential, PTF = Potentiated Twitch Force, RTF = Resting Twitch Force, TB = Triceps Brachii, VMA = Voluntary Muscle Activation

### Reliability

With the exception of moderate internal consistency (0.7) for biceps brachii’s M-wave, and acceptable (0.7 ≤ α < 0.8) reliability for resting twitch force, ICC reliability scores were excellent (0.82 ≤ α < 0.94) for MVC forces, potentiated twitch forces, biceps and triceps brachii EMG (Table 2).

**Table 2.**
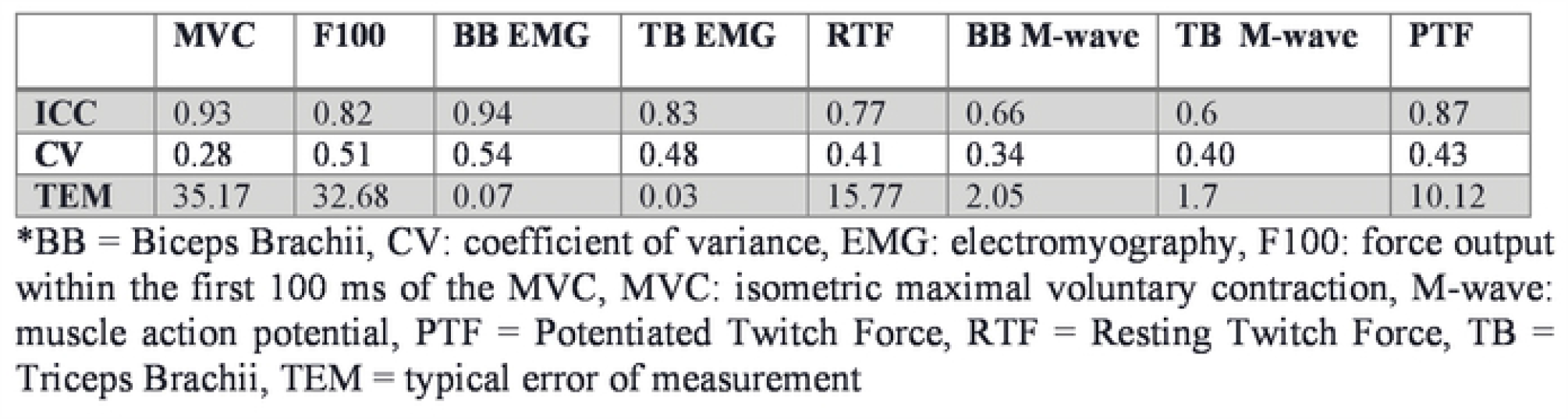
Daily intraclass correlation coefficient reliability (initial resting position test performed on separate days).

#### Voluntary contractile properties

##### Elbow flexors MVC

A significant seated position x time interaction (*F*_(2,16)_ = 5.07, *p*=0.02, 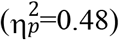 (Figure 3 was observed for elbow flexor MVC force. The interaction revealed 9.1% and 21.1% MVC force decreases (initial>pre-fatigue) for upright and inverted positions, respectively. Furthermore, there were 21.4% and 14.2% force decreases from the initial and pre-test to post-test respectively for the upright position. Similarly, there were 33.1% and 17.7% decrements from the initial and pre-test to post-test respectively for the inverted position. A non-significant, medium effect size, seated position x BFR interaction (*F*_(1,8)_ = 3.79, *p*=0.08, 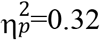); demonstrated 9.1% lower MVC forces with BFR compared to without BFR for inverted seated positions.

**Figure 3:**
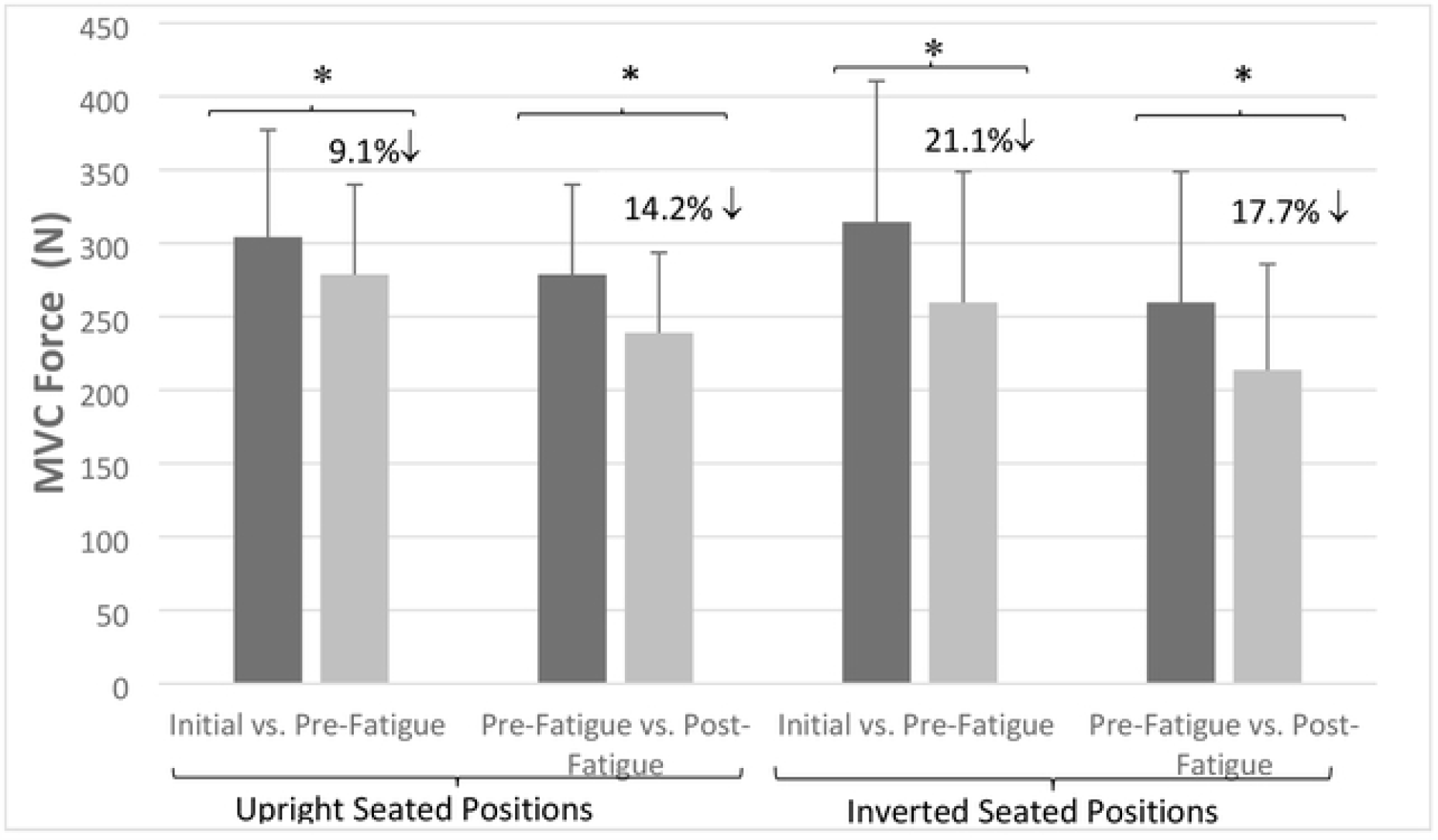
Isometric maximal voluntary contraction force (MVC) interaction effects for seated position and time. Star(*) symbol represents that significant MVC force decreased between initial and pre-fatigue tests, for upright and inverted seated positions. Means and standard deviations are illustrated. There was no statistically significant BFR interaction.

### Voluntary muscle activation (%)

A non-significant, but large eta^2^ magnitude, BFR x time interaction revealed 7.9% and 16.4% decreases post-30-s MVC (*F*_(1,4)_ = 5.48, *p*=0.07, 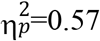) for without BFR and BFR conditions, respectively (Figure 4).

**Figure 4:**
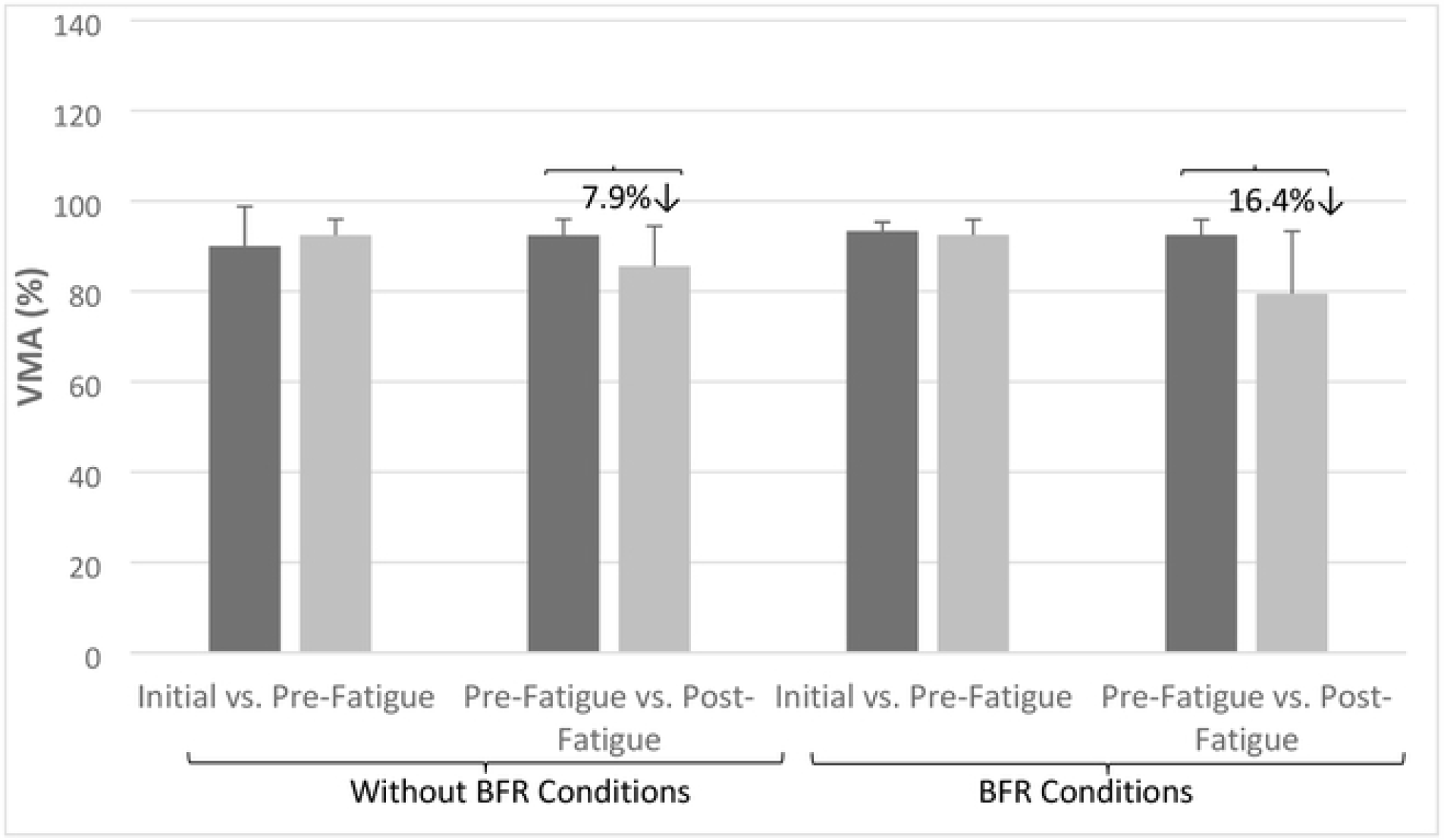
Voluntary muscle activation (VMA) interaction effects for blood flow restriction (BFR) and time. There was a non-significant effect for percentage of VMA (*p*=0.07), with 7.9% and 16.4% decreases post-fatigue for without BFR and BFR conditions. Means and standard deviations are illustrated. There was no statistically significant interaction between seated positions (inverted versus upright).

### Evoked twitch contractile properties

A significant seated position x time interaction (*F*_(1,11)_ = 5.80, *p*=0.03, 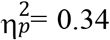), for resting pre-MVC twitch force revealed that initial values exceeded pre-30-s MVC, by 29.4% for inverted seated positions (Figure 5). No significant effects or interaction were found for baseline time to peak twitch force. Meanwhile, a non-significant, large magnitude effect size effect was observed (*F*_(1,6)_ = 4.81, *p*=0.07, 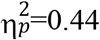), in terms of seated position and BFR, when comparing upright values (without BFR<BFR, 3.9%↑) with inverted seated positions (without BFR>BFR, 5.6%↓). No significant effects were observed for half relaxation time-twitch force.

**Figure 5:**
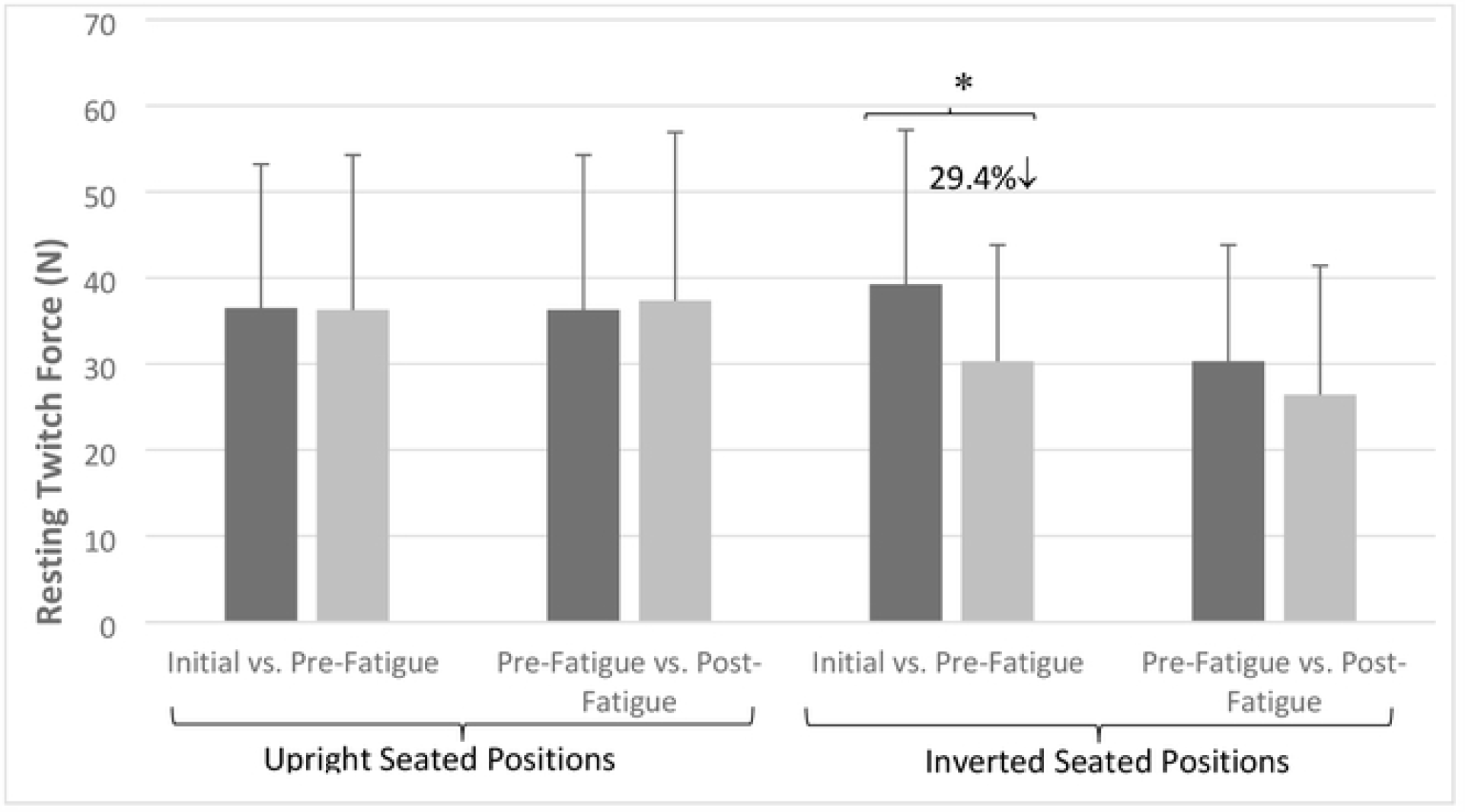
Resting twitch force interaction effects for seated position and time. Star (*) symbol represents significant decreases between initial and pre-fatigue tests, for upright and inverted seated positions. Means and standard deviations are illustrated. There was no statistically significant BFR interaction.

### Biceps and triceps brachii M-wave

A significant seated position x BFR interaction (*F*_(1,7)_ = 5.69, *p*=0.04, 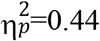), revealed that the without BFR condition exceeded BFR condition biceps brachii’s M-wave by 30.4% and 12.5% for upright, and inverted seated positions, respectively (Figure 6).

**Figure 6:**
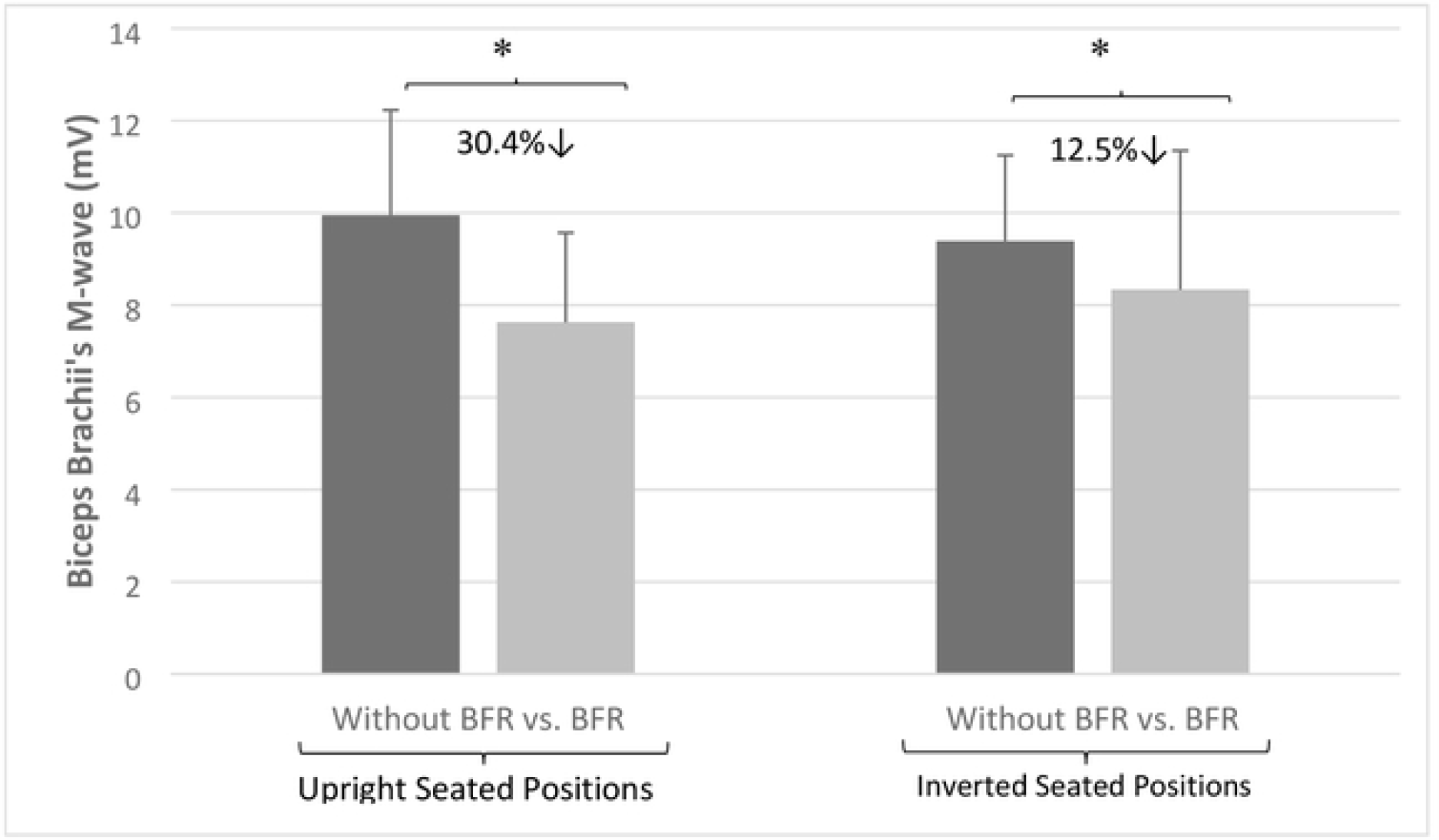
Biceps brachii M-wave interaction effects for seated position and BFR. Star(*) symbol represents significant decreases in amplitude of M-wave for biceps brachii, between without BFR and BFR, with 30.4% and 12.5% for upright and inverted seated positions, respectively. BFR = blood flow restriction. Means and standard deviations are illustrated.

### Evoked potentiated twitch contractile properties

Significant seated position x time interaction (*F*_(2,10)_ = 7.91, *p*=0.009, 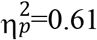) was observed for potentiated twitch force (PTF). The pre- and post-30-s MVC results showed 65.3% and 75% PTF decreases for upright and inverted positions from pre-to post 30-s MVC, respectively (Figure 7).

**Figure 7:**
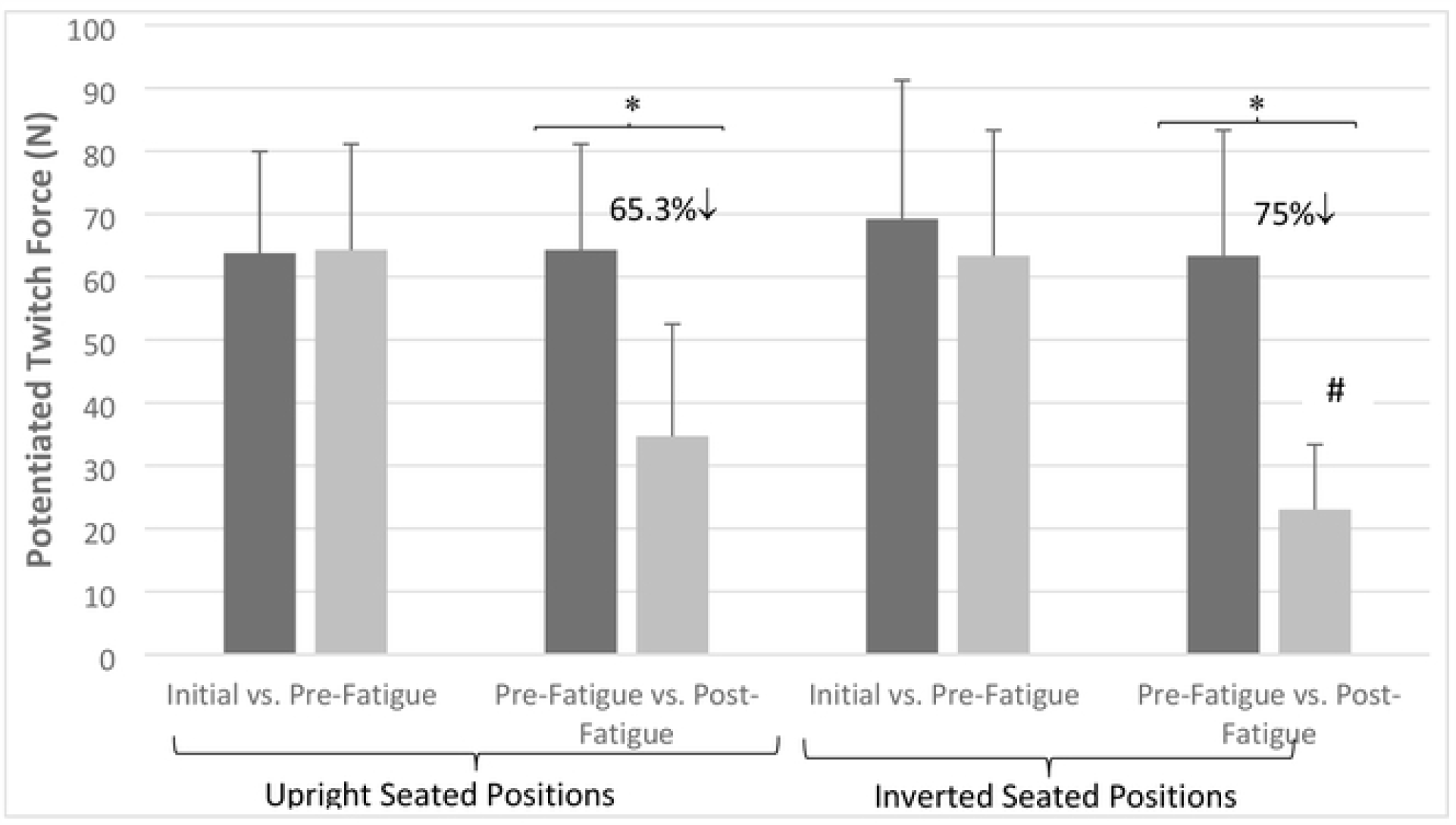
Potentiation twitch force (PTF) interaction effects for seated position and time. Star(*) symbol represents that significant potentiation twitch force decreases between pre- and post-fatigue tests, for upright and inverted seated positions. The hashtag or number symbol(#) indicates that PTF tested with inversion post-fatigue was significantly lower than all other times and conditions. Means and standard deviations are illustrated. There was no statistically significant BFR interaction.

### Fatigue – force relation

When comparing force output values, at 0-5 s and 25-30 s intervals of the 30-s MVC task in the without BFR conditions versus 0-5 s and 25-30 s of BFR conditions, there was 4.1% force decreases for BFR effects (*F*_(1,11)_ = 5.54, *p*=0.03, 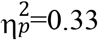).

### Fatigue - EMG relation

### Biceps brachii

Furthermore, contrasts revealed a non-significant, medium magnitude effect size, seated position x time interaction (*F*_(1,10)_ = 3.84, *p*=0.07, 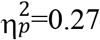); with 45.6% and 38.6% decreases in biceps brachii EMG median frequency during upright without BFR and BFR conditions respectively, versus similar 32.8% and 41.4% median frequency decreases during the inverted seated position for without BFR and BFR conditions, respectively (Table 2).

### Perceived pain - pain scale

Seated position x BFR interaction revealed significant differences (*F*_(1,12)_ =6.55, *p*=0.02, 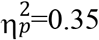) between upright (without BFR<BFR, 0.64 to 1.53, 58.1%↑) and inversion (without BFR>BFR, 2.70 to 2.43, −11.1%) positions. For the interaction of BFR x time (*F*_(2.05,24.62)_ = 4.68, *p*=0.01,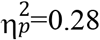), a 4.6% pain scale decrease between 20-40 s and 40-60s intervals for the without BFR condition, contrasted with an overall (both without BFR and BFR) increase from initial (upright/inverted) to post-30-s MVC measures. Furthermore, a seated position x BFR x time interaction (*F*_(2.87,34.47)_ = 3.18, *p*=0.03,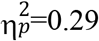) showed significant differences when comparing without BFR and BFR, upright versus inverted conditions between 0-20 s and 20-40 s intervals (−32.6% (0.61 to 0.46) without BFR and 13.6%↑ (1.46 to 1.69) for BFR upright positions, versus 7.8%↑ (2.69 to 2.92) and 3.2%↑ (2.38 to 2.46) for inverted without BFR and BFR conditions), and between 40-60 s and post-30-s MVC (77.5%↑ (0.38 to 1.69)) and 35.2%↑ (1.84 to 2.84) for upright without BFR and BFR versus 13.9%↑ (2.84 to 3.30) and 26%↑ (2.61 to 3.53) for inverted without BFR and BFR conditions).

A main effect for time (*F*_(1.37, 16.49)_ = 11.39, *p*=0.002,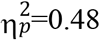) showed 34.6% (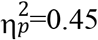, initial<pre-30-s MVC), 14% (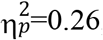, pre-30-s MVC<0-20 s interval), and 32.3% (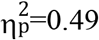, 40-60 s interval<post-30-s MVC) pain scale increases. Main effects for seated position and BFR showed 26.5% (upright<inversion) (*F*_(1,12)_ = 14.56, *p*=0.002,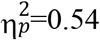) and 15.6% increases (without BFR<BFR) (*F*_(1,12)_ = 7.85, *p*=0.01,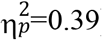) for pain scale.

## DISCUSSION

Prior studies have reported upon the deficits related to global (whole body) effects of inversion-induced changes in hydrostatic pressure (Hearn et al. 2009; Johar et al. 2013; Neary et al. 2015; Paddock et al. 2009; Smith et al. 2013). Similarly, there is a body of literature exhibiting impairments associated with local (muscle group) BFR-induced increases in perfusion pressure (Amann et al. 2008, Fitzpatrick et al. 1996, Hobbs and McCloskey 1987, Sundberg and Kaisjer 1992). The present study is the first to investigate the separate and combined effects of inversion (increased hydrostatic pressure) and BFR (increased perfusion pressure) on voluntary and evoked contractile properties. Major findings of this study were that inversion induced significantly greater decreases in resting twitch and elbow flexors MVC forces before the 30-s MVC task. Following the 30-s MVC task, inversion induced greater decreases in PTF. BFR led to overall (inversion and upright) detriments in MVC force, as well as greater decreases than without BFR while in the inverted position. Furthermore, there was a greater decrease in M-wave amplitude for the upright versus inverted position. In addition, the concomitant application of inversion of the participant with BFR of the elbow flexors only amplified neuromuscular performance impairments to a small degree. Representative traces of the measures are presented in Figure 8.

**Figure 8:**
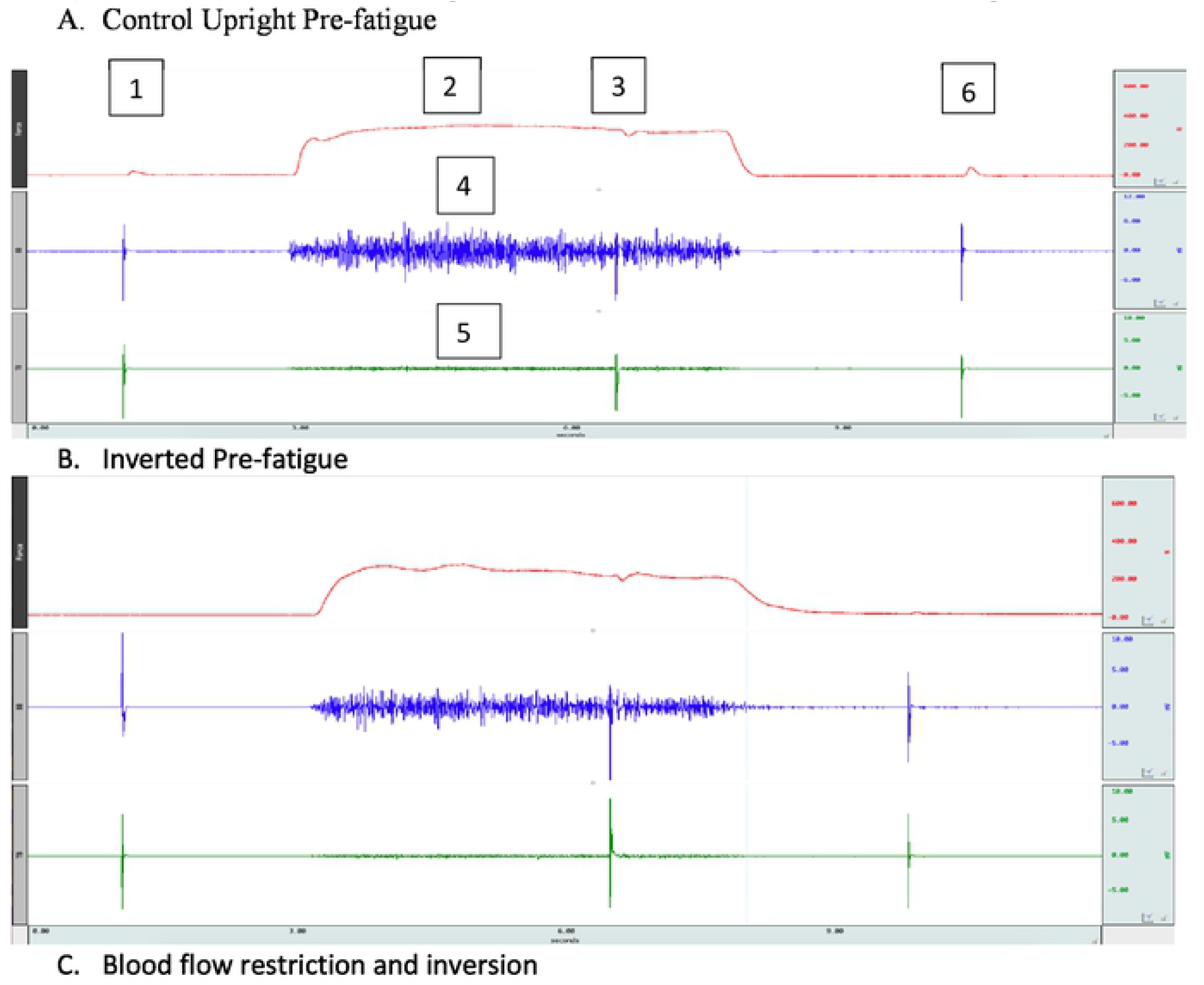
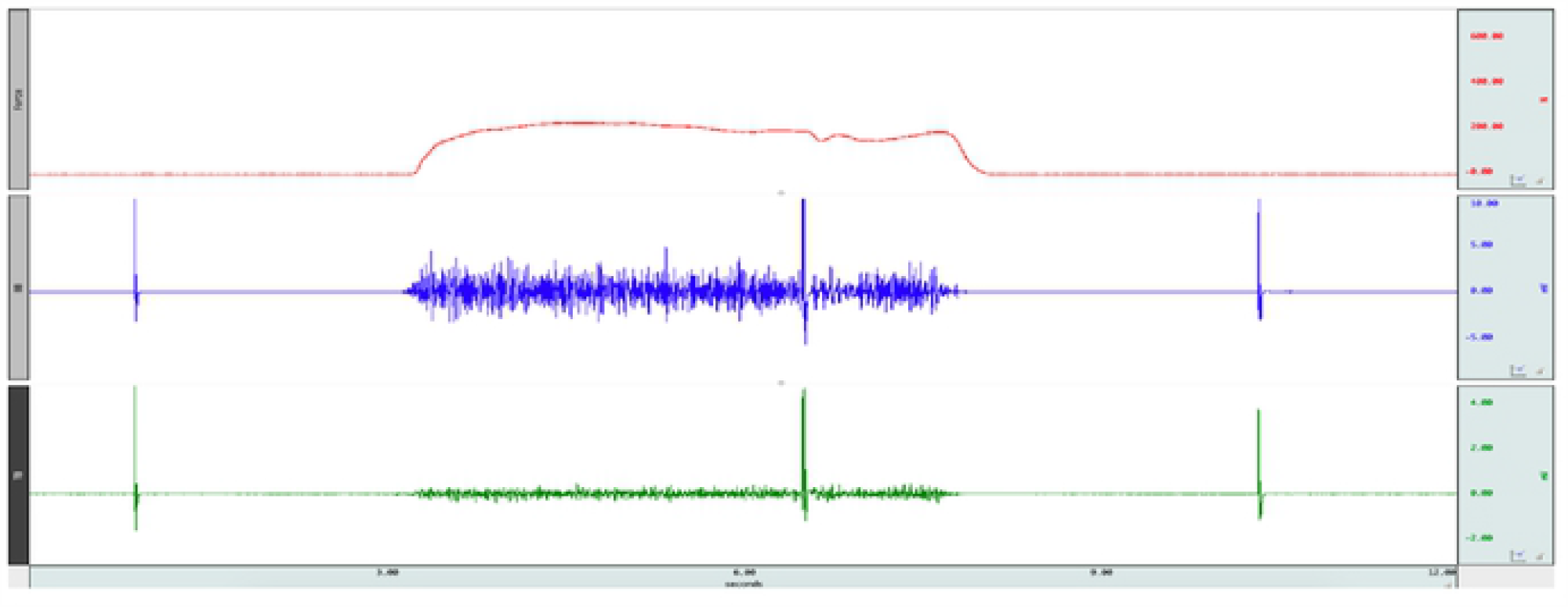
Representative 1) evoked resting twitch, 2) MVC force, 3) interpolated twitch technique (ITT), 4) biceps brachii EMG, 5) triceps brachii EMG and 6) potentiated twitch force (PTF) tracings presented under different conditions/posture. Top channel represents elbow flexor MVC force, second channel illustrates biceps brachii EMG and third channel shows triceps brachii EMG.

### Inversion Effects

#### Evoked contractile properties

Significant reductions in pre-test evoked twitch force with inversion are partially in accord with a previous inversion study that showed non-significant, but moderate magnitude decreases (18.6%, effect size [ES] = 0.74) for elbow flexors resting twitch force from upright to the inverted position contrasting with non-significant moderate magnitude increases (17.5%, ES = 0.76) for knee extensors (Neary et al. 2015). Similarly, greater 30-s MVC-induced PTF decreases with inversion versus upright, in this research, also conforms with Neary et al. (2015), who showed non-significant (*p*=0.06), small magnitude, decreases for elbow flexor PTF (11.8%, ES = 0.44); whereas they reported large magnitude increases (*p*=0.03, ES = 1.27, 27.3%) for leg extensor PTF forces during seated inversion.

With inversion, the position of the arm and leg induces higher and lower hydrostatic pressure respectively, hence contributing to the contrasting arm and leg responses in the Neary et al. (2015) study. With high gravitational pressure, previous animal studies have shown an altered function of acetylcholine receptors (Heinemann et al. 1987), relative decrease in psoas single muscle fibre isometric force (Geeves and Ranatunga 1987), decreased tetanic force for extensor digitorum longus muscle (Ranatunga and Geeves 1991), and decreased Ca^++^ influx into the nerve terminal (Grossman and Kendig 1990). In humans, Sundberg and Kaisjer (1992) showed that a graded occluded lower limb (up to 50 mmHg to increase pressure) under upright conditions caused a progressive decrease in muscle function. These results suggest higher hydrostatic or perfusion pressures on the arm during inversion can significantly reduce evoked forces.

In contrast, Paddock and Behm (2009) reported no significant alterations to evoked twitch properties with seated inversion. In the Paddock and Behm study, only male subjects were recruited, and they spent 30-s rather than 1-min in the inverted position. In addition, after each contraction, the participants were returned to an upright position for 2-minutes before adopting the inverted position again for another series of voluntary and evoked contractions. Hence, the differences in duration and recovery from inversion may have attenuated changes in hydrostatic pressure while sex differences may have contributed to the differences in evoked twitch property results.

#### Voluntary contractile properties

The 21.1% inversion-induced MVC force decrease in the present study aligns with the 18.9-21.1%, 10.4%, and 6.1% impairments reported by Johar et al. (2013), Hearn et al. (2009) and Paddock and Behm (2009), respectively. It might be argued that the relatively less substantial force decreases experienced by the participants in the Hearn et al. (2009) and Paddock and Behm (2009) studies may be attributed to shorter durations of inversion (30-s or less in each study) with 2-min recovery periods to an upright position between each MVC. However, Johar and colleagues had very similar force decreases as the present study and had participants exert MVCs for 6-s following rapid rotations of 1- or 3-s to an inverted position. Thus MVCs were competed in less than 10-s in the Johar et al. (2013) study. Neary et al. (2015) reported no significant change in elbow flexor MVC force, which might be attributed to the highly trained track and field (athletics) athletes recruited in their study, who were permitted 1-min recovery periods between contractions. Hence, the relatively lower or non-significant force decrements in the Hearn et al. (2009), Paddock and Behm (2009) and Neary et al. (2015) studies respectively may be more related to the greater recovery periods between inversion and return to upright positions.

The decreases in elbow flexors MVC force with inversion can be related to several factors. The decreased force output can be related to a muscle stiffening mechanism, associated with a threat of instability (Carpenter et al. 2001), that the participants could have perceived when inverted with only straps holding them in the chair. Adkin et al. (2002) found that the stiffening strategy can negatively influence voluntary movement. Increased co-contractile activity with inversion has been previously reported as a factor counteracting target force output (Hearn et al. 2009; Paddock and Behm 2009), however there were no significant increases in triceps brachii EMG in the present study. It is possible that an increased focus on stabilizing functions of the shoulders, and trunk muscles (Arokoski et al. 2001), could negatively impact the force output of the elbow flexors (Anderson and Behm 2004; Drinkwater et al. 2007). A shift from mobilizing to stabilizing strategies of the neuromuscular system has been reported to contribute to force reduction (Anderson and Behm 2004, Behm and Anderson 2006).

There were no significant inversion-induced decrements in biceps brachii EMG, which contrasts with 21.7%-35.9%, 26.6%, 47.9% decreases with Johar et al. (2013), Paddock and Behm (2009) and Hearn et al. (2009) studies, respectively. As mentioned previously, these contrasts may be attributed to the greater inversion and contraction recovery periods or recruited population (i.e., track and field athletes) in the cited studies. Hence, while these other studies postulated inversion-induced neural inhibition due to increases in cerebral blood pooling and intracranial pressure (Bosone et al. 2004; Matsubara et al. 1990; Warkentin et al. 1992), or attenuated sympathetic drive (Bosone et al. 2004; Cooke et al. 2004; Cooke and Dowlyn 2000; Fitzpatrick et al. 1996), the lack of EMG and ITT changes in the present study suggests that neural influences did not play a major role in MVC force reductions. Voluntary activation (ITT) was decreased following the 30-s MVC for both BFR and no-BFR but EMG did not significantly change. The curvilinear nature of the EMG-force relationship often resulting in a plateau at high voluntary forces diminishes the EMG sensitivity to force changes at high contraction intensities (Behm and St-Pierre 1997b).

### Blood Flow Restriction (BFR)

#### Voluntary and evoked contractile properties

Overall, BFR induced 4.8% lower MVC force values than without BFR as well as 9.1% lower MVC forces with BFR in an inverted position versus without BFR. This relative impairment is similar to the BFR-induced deficits for time to peak twitch. Without BFR, time to peak twitch increased 3.9% versus BFR when upright versus a 5.6% decrease with BFR vs. without BFR when inverted. Overall, BFR also impaired M-wave amplitudes by 21% (30.4% and 12.5% deficits for upright, and inverted seated positions respectively) compared to without BFR.

These findings are similar to the force (Sundberg and Kaisjer 1992) and neuromuscular efficiency deficits reported by Hobbs and McCloskey (Hobbs and McCloskey 1987) with graded ischaemia. Ischaemia and the associated pain can contribute to central nervous system inhibition (Bigland-Ritchie et al. 1986; Garland 1991) adversely affecting force output. Similar to inversion, the occlusion of venous blood flow return would increase hydrostatic / perfusion pressure at the muscle resulting in similar force deficits seen in animals (Geeves and Ranatunga 1987; Ranatunga and Geeves 1991) and humans (Sundberg and Kaisjer 1992). Hogan et al. (1994) reported force decrements with diminished oxygen delivery to working muscle. Copithorne et al. (2020) showed that BFR induced more significant decreases (∼80%) in time to task failure during a sustained isometric fatiguing task, in comparison with normal blood flow condition. Furthermore, impairments in the evoked time to peak twitch and M-wave could be partly attributed to an alteration in muscle fibre conduction velocity (Farina et al. 2004) with nerve compression (Ogata et al. 1991) as well as impaired sarcoplasmic reticulum Ca^2+^ uptake (Place et al. 2009). With median nerve compression in the carpal tunnel, Lunderborg et al. (1982) observed a change in endoneural microcirculation. While it would be logical to suspect peripheral alterations with BFR, central responses can also be influenced.

The greater BFR-induced decrease (*p*=0.07, 16.4%) in biceps brachii EMG versus without BFR (7.9%) may reflect a greater hypoxia placing a greater emphasis on anaerobic metabolism, resulting in an earlier higher threshold motor unit recruitment accelerating the onset of force decrements (Gerdle and Fugl-Meyer 1992; Moore et al. 2004). Occluding arterial flow in the arm, during the post-exercise period, Bigland-Ritchie et al. (1986) observed an impairment of neuromuscular transmission.

#### Perceived pain

Previous seated inversion studies (Johar et al. 2013, Neary et al. 2015; Paddock and Behm 2009) have mentioned that there was a sense of discomfort (i.e., distinct swelling around the head region) during inversion but did not directly measure or report the degree of discomfort. Overall, pain perception tended to be higher with BFR (15.6%) as well as when inverted (26.5%). Increased pain perception with inversion would be related to the increased hydrostatic pressure around the head region (Paddock and Behm 2009). BFR would induce a hypoxic muscle environment resulting in a greater reliance on anaerobic metabolism, higher accumulation of metabolites and with the BFR, an inability to dispose these metabolites (Hogan et al. 1994; Sundberg and Kaisjer 1992). A compressed limb can lead to ischaemia accumulating metabolites and with increased local acidity would activate group III and IV pain afferents promoting a sensation of increased discomfort and pain (Hollander et al. 2010; Leonard et al. 1994; Marchettini et al. 1996).

#### Inversion and BFR Interactions

The data tended to indicate that BFR with inversion induced small (∼5-12.5%) multiplicative effects upon voluntary and evoked muscle force and activation. A non-significant (p=0.07) but large partial eta^2^ magnitude BFR x inversion interaction showed that evoked twitch forces had a BFR-induced greater increase (5.6%) than without BFR in the inverted position. In contrast, MVC forces with BFR showed greater decreases (−9.1%) than without BFR for the inverted seated position. The M-wave amplitude was more impaired by BFR (−12.5%) than without BFR in the inverted position. Finally, pain perception increased with BFR (11.1%) versus without BFR with the inverted position. Thus, the combination of inversion and BFR provided some amplification of the deficits compared to inversion alone (i.e., decreased elbow flexor resting twitch and MVC forces). The BFR-induced hypoxic environment attenuates blood substrates leading to a greater accumulation of metabolic by-products that could negatively impact force production. On the other hand, BFR may increase neuromuscular excitation (i.e., increased M-wave amplitudes). Neural compression and blood flow impairment have elevated EMG with submaximal contractions (Karabulut et al. 2010; Yasuda et al. 2006). Copithorne et al. (2020) reported that blood flow occlusion led to a more rapid and greater increase in motoneuron excitability but had no effect on motor cortical excitability suggesting that group III/IV afferent feedback was the primary cause of the enhanced motoneuron excitability. Thus, BFR can induce both peripheral impairments and central excitation. These contrasting effects in combination with inversion-induced (i.e., increased hydrostatic pressure) alterations provided small but not major amplification of the deficits.

There were some minor limitations to the research. Due to the research procedure, data could only be obtained from the right limb. While the partial BFR was set relative to the position (upright/inverted), it could not exclusively indicate 50% BFR upright was exactly the same as inverted.

## Conclusions

Hydrostatic pressure-induced peripheral muscle impairments with the inverted position, may have contributed to decreases in elbow flexor twitch, PTF, and MVC forces before the fatigue task. BFR did decrease voluntary force and M-wave amplitudes with only a small additional increase to the inversion-induced impediments. A decrease in force or power output, with an increased perceived difficulty in stressful and life-threatening conditions such as accidents involving inversion or blood flow restriction (i.e., overturned vehicles, aerial maneuvers) can be life threatening. The results provide insights into the separate and combined effects of global (inversion) and local (BFR) increases in hydrostatic/perfusion pressure on voluntary and evoked contractile properties.

## AUTHOR CONTRIBUTIONS

HA and DGB conceived the research question and protocol. HA, NH, and SA were involved with data collection and analysis. HA wrote the first draft of the manuscript. DGB, DCB, and UG reviewed and provided extensive revisions to the paper.

## ABBREVIATIONS

BFR: Blood flow restriction
EMG: electromyography
ICC: intraclass correlation coefficient
ITT: interpolated twitch technique
MVC: maximal voluntary contraction
PTF: potentiated twitch force

